# Genome-scale TraDIS reveals dynamic and conserved fitness requirements of *Salmonella* Typhimurium across sequential host niches during porcine infection

**DOI:** 10.64898/2026.03.03.709262

**Authors:** Antonio Romero-Guillén, Joaquín Bernal-Bayard, Lars Barquist, Francisco Ramos-Morales, Sara Zaldívar-López, Tránsito García-García, Juan J. Garrido

**Affiliations:** Departamento de Genética, Grupo de Inmunogenómica y Patogénesis Molecular, UIC Zoonosis y Enfermedades Emergentes ENZOEM, Universidad de Córdoba, Córdoba, España; Instituto Maimónides de Investigación Biomédica de Córdoba (IMIBIC), Grupo GA-14, Córdoba, España; Departamento de Genética, Facultad de Biología, Universidad de Sevilla, Sevilla, España; Helmholtz Institute for RNA-based Infection Research (HIRI), Helmholtz Centre for Infection Research (HZI), Würzburg, Germany; Faculty of Medicine, University of Würzburg, Würzburg, Germany; Department of Biology, University of Toronto, Mississauga, Ontario, Canada; Department of Cell and Systems Biology, University of Toronto, Toronto, Ontario, Canada

## Abstract

Non-typhoidal salmonellosis remains a major global cause of foodborne gastrointestinal disease, with pigs representing an important reservoir of *Salmonella enterica* serovar Typhimurium. The emergence of host-adapted, multidrug-resistant lineages has further reinforced the need to understand the genetic basis of bacterial persistence and pathogenicity within physiologically relevant hosts.

Here, we applied transposon-directed insertion sequencing (TraDIS) to systematically define the genetic requirements of a highly host-adapted, multidrug-resistant *S*. Typhimurium DT104 isolate across sequential host niches during porcine infection. A high-density transposon mutant library (∼1.2 million mutants) was subjected to *in vivo* selection in the ileal mucosa and mesenteric lymph nodes, as well as *ex vivo* infection of primary porcine neutrophils.

Across all conditions, we identified 1,813 conditionally essential genes, revealing strong niche-specific fitness signatures and a progressive increase in selective stringency along the infection route. Ileal colonization was primarily driven by determinants of invasion, motility and lipopolysaccharide biosynthesis. In contrast, survival within neutrophils depended on resistance to antimicrobial stresses and extensive metabolic rewiring, whereas persistence in lymph nodes required a broader functional repertoire integrating virulence, motility, envelope remodelling and metabolic adaptation, including bacterial microcompartment-associated pathways.

Despite this marked heterogeneity, we identified a conserved porcine host-conditioned essential genome, comprising core invasion functions, RNA metabolism and genome maintenance pathways, and central metabolism. Notably, the outer membrane lipid asymmetry system Mla and the twin-arginine translocation (Tat) pathway emerged as conserved bottlenecks for *in vivo* fitness across all host-associated environments.

Together, these findings establish a hierarchical model of *S.* Typhimurium adaptation during porcine infection and provide a systems-level view of tissue-specific and conserved genetic requirements underpinning persistence in a major zoonotic reservoir.

## Introduction

Non-typhoidal salmonellosis remains one of the leading causes of foodborne gastrointestinal disease worldwide, posing a substantial public health burden in both industrialized and developing countries ^1^. Among food-producing animals, pigs constitute a major reservoir of *Salmonella enterica* serovar Typhimurium, facilitating zoonotic transmission through the food chain and contributing to the persistence of endemic lineages in agricultural settings^2^. Of particular concern is the emergence of host-adapted, multidrug-resistant *S.* Typhimurium strains, such as phage type DT104 and the monophasic variant 1,4,[5],12:i:-, which compromise therapeutic efficacy and underscore the urgent need to better understand the bacterial determinants that underpin persistence and pathogenicity within the porcine host ^3^.

Following oral acquisition, *S.* Typhimurium undertakes a complex and highly structured infection route in pigs. Initial colonization occurs in the distal ileum, where the bacterium must overcome multiple barriers including intestinal mucus, resident microbiota, antimicrobial peptides, and epithelial innate immune defenses^4^. Successful invasion of the intestinal epithelium enables subsequent dissemination to lymphoid tissues, particularly mesenteric lymph nodes, which represent a critical bottleneck for systemic persistence^5^. In parallel, *Salmonella* encounters professional phagocytes such as neutrophils, which impose intense antimicrobial pressure through oxidative burst, antimicrobial peptides, and nutrient deprivation^6^. Each of these anatomical niches imposes distinct physiological constraints, requiring precise and dynamic adaptation of bacterial virulence programs, metabolism, and envelope integrity.

Much of the current understanding of *Salmonella* pathogenesis has been derived from murine models, which have been instrumental in elucidating conserved mechanisms of invasion, intracellular survival, and immune modulation^7^. However, important differences exist between mice and pigs in gastrointestinal physiology, immune architecture, microbiota composition, and infection outcome. In pigs, *S.* Typhimurium often establishes subclinical or persistent infections that closely resemble non-typhoidal salmonellosis in humans, positioning the porcine host as a highly relevant translational model^8^.

Transposon-directed insertion sequencing (TraDIS) has emerged as a powerful approach to systematically map gene essentiality and fitness landscapes at genome scale, enabling the identification of conditionally essential genes required for survival under specific environmental or host-imposed stresses^9,10^. Applied to enteric pathogens, TraDIS has revealed key virulence determinants, metabolic pathways and stress response systems required during infection^11^. Previous attempts to comprehensively assign the role of Salmonella genes relied largely on parenteral infection of atypically susceptible mice and therefore did not reflect the genetic requirements of *Salmonella* during intestinal colonization of food-producing animals infected via the natural oral route. To identify genes relevant to colonization of animal reservoirs and, consequently, zoonotic transmission, genome-scale TraDIS analyses have since been applied to food-producing animals, including chickens, pigs and calves, following oral infection^12^. However, these studies have largely examined individual intestinal compartments in isolation, providing limited insight into how bacterial genetic requirements are reshaped across sequential host niches encountered during natural infection. In particular, how selective pressures shift as *Salmonella* transitions from the intestinal mucosa to innate immune cells and lymphoid tissues within the same host remains poorly understood.

Here, we address this gap by performing a genome-wide TraDIS analysis of a highly host-adapted, multidrug-resistant *S.* Typhimurium DT104 isolate during infection of pigs. By interrogating bacterial fitness in three biologically and temporally linked environments—the ileum, primary porcine neutrophils, and mesenteric lymph nodes (MLN) —we provide a systems-level view of how *Salmonella* reshapes its genetic requirements along the porcine infection route (**Figure 1A**). This multicontext approach enables us to distinguish tissue-specific fitness determinants from a conserved host-conditioned essential genome that underpins survival across diverse host niches.

**Figure 1.**
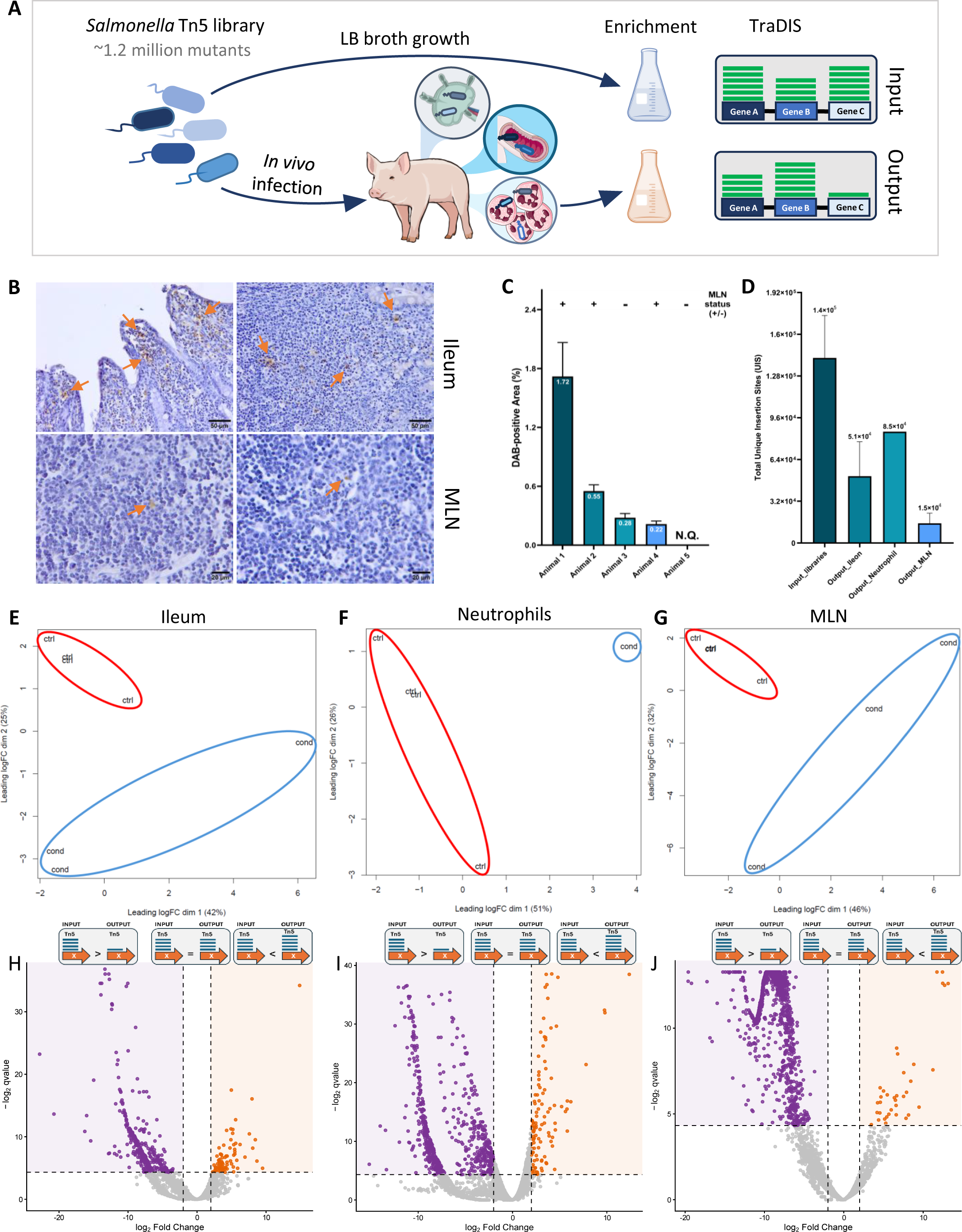
Genome-wide identification of *Salmonella* genes essential for invasion and colonization in the porcine intestinal model. **A.** Schematic of the protocol used for the Salmonella TraDIS screen in pigs. **B**. Detection of Salmonella in porcine tissues by immunohistochemistry; DAB-positive staining is indicated by orange arrows. **C.** Quantification of DAB-positive area (%) in ileal samples (mean ± SD); bacterial presence in mesenteric lymph nodes (MLN) was assessed qualitatively by immunohistochemistry. **D**. Average number of unique transposon insertion sites (UIS) detected per condition across sequenced TraDIS libraries (mean ± SD). **E-G.** MDS-plot showing clustering of input and output libraries based on gene-level fitness profiles. **H-J**. Volcano plot illustrating differential mutant abundance between infected and control conditions, dashed lines indicate significance thresholds (log2 fold change ±2 and q-value ≤ 0.05).

Our analysis reveals striking tissue-specific genetic signatures reflecting niche-adapted strategies, including invasion and motility in the intestinal mucosa, stress resistance and metabolic rewiring in neutrophils, and extensive virulence and metabolic integration within lymphoid tissue. At the same time, we identify a compact set of conserved fitness determinants required across all porcine infection environments, highlighting fundamental physiological bottlenecks imposed by the host. Notably, pathways involved in outer membrane lipid asymmetry maintenance and twin-arginine–dependent protein export emerge as universally essential for *in vivo* fitness, revealing previously underappreciated vulnerabilities in *S.* Typhimurium host adaptation.

Together, this work establishes a comprehensive genetic atlas of *S.* Typhimurium fitness across key porcine infection niches and provides mechanistic insight into how host-imposed constraints shape pathogen survival in a large-animal model. By integrating tissue-specific and conserved genetic requirements, our findings advance understanding of *Salmonella* pathogenesis in pigs and identify candidate pathways for targeted intervention to control non-typhoidal salmonellosis at the animal–human interface.

## Material and methods

### Bacterial strain and Tn5 library generation

The bacterial strain employed in this study was *Salmonella enterica* subsp. *enterica* serovar Typhimurium phage type DT104 SP11. This strain is an environmental isolate recovered from a carrier pig in a porcine farm in Leon, Spain ^13^. Prior to library construction, the antibiotic susceptibility profile of the strain was determined to guide downstream selection conditions. Minimal inhibitory concentrations (MICs) for ampicillin, chloramphenicol, kanamycin and gentamicin were determined in Luria–Bertani (LB) broth using two-fold serial dilutions (**Figure S1A**). Cultures were incubated at 37 °C with shaking for 24 h in 96-well plates, and bacterial growth was monitored hourly by measuring optical density at 600 nm (OD_600_) using a Varioskan™ LUX microplate reader.

Tn5 mutant library of the *S.* Typhimurium strain was generated using the mini-Tn5 carrier plasmid pBAM1, that has an ampicillin resistance gene inserted in its backbone (plasmid marker) and a kanamycin resistance gene inserted in the transposable region (**Figure S1B**)^14^. Plasmid transfer was performed by conjugation following a protocol adapted from Karash *et* al. ^15^. Briefly, the donor strain *E. coli* S17-1λPIR was transformed with pBAM1 by heat-shock and transformants were selected on LB agar supplemented with ampicillin (100 µg/ml) and kanamycin (50 µg/ml). Overnight cultures of both donor (*E. coli* S17-1λPIR pBAM1) and recipient (*S.* Typhimurium DT104 SP11) strains were washed once with 10 mM MgSO₄ and once with phosphate-buffered saline (PBS, 1x). Equal volumes of donor and recipient cultures were mixed and plated onto LB agar without antibiotics, followed by incubation for 5 h at 37 °C to allow conjugation. After incubation, bacterial biomass was recovered using a sterile inoculation loop and resuspended in 1 ml of 10 mM MgSO₄. Ten-fold serial dilutions were plated onto LB agar supplemented with kanamycin (50 µg/ml) and chloramphenicol (30 µg/ml) to selectively recover *S.* Typhimurium transposon mutants. Donor and recipient strains plated independently under the same selective conditions served as negative controls for conjugation (**Figure S1C**). In parallel, the remaining conjugation mixture was plated undiluted onto selective LB agar to obtain the full transposon mutant library. After overnight incubation at 37 °C, colonies were harvested and pooled to generate the final Tn5 mutant library. Aliquots were stored at −80 °C in LB broth supplemented with kanamycin and 7% (v/v) DMSO.

The Tn5 mutant library was validated for the presence of residual pBAM1 plasmid, since the intrinsic ampicillin resistance of *S.* Typhimurium DT104 SP11 prevented plasmid counter-selection on ampicillin-containing media. Therefore, plasmid persistence was assessed by colony PCR using primers targeting the transposase A gene (*tnpA*) sequence of the pBAM1 backbone (**Figure S1D**). A total of 60 randomly selected mutants from independent conjugations were screened for the presence of *tnpA*. Randomness of Tn5 insertion was evaluated in 20 randomly selected mutants using a semi-random nested PCR strategy as described by de la Rosa-Altura et al.^16^ (Figure S1F). Briefly, initial amplification was performed using an arbitrary primer (ST-ACGCC) in combination with either ME-I-extR (upstream end) or ME-O-extF (downstream end) of the transposon. A second round of PCR employed a nested arbitrary primer (ST1) together with ME-I-intR or ME-O-intF, respectively. Random insertion of Tn5 is expected to generate distinct PCR profiles across mutants, which were confirmed by agarose gel electrophoresis. Selected amplification products were excised from gels, purified using the Wizard® SV Gel and PCR Clean-Up System (Promega), and submitted for Sanger sequencing (Stab Vida) using primer ST1. Insertion sites were then mapped against the S. Typhimurium DT104 SP11 reference genome using SnapGene Viewer v7.2.1.

### Infection model and recovery of *in vivo* Tn5 mutant libraries

The Tn5 mutant library was refreshed by thawing the bacterial stock at 37 °C followed by incubation in LB broth supplemented with kanamycin (1:10 dilution) for 1 h with shaking. Cells were then washed once in PBS and inoculated into fresh LB broth with kanamycin at a final 1:50 dilution. Cultures were grown at 37 °C with agitation to mid-exponential phase (OD₆₀₀ ≈ 0.8) prior to infection. Five crossbred weaned piglets (male and female), approximately 4 weeks of age, were used in this study. All animals were confirmed negative for *Salmonella* by fecal culture prior to infection. Animal procedures were performed at the University of León (Spain) following a previously described experimental model^4^. Piglets were orally inoculated with 6.25 mL of activated Tn5 mutant library containing approximately 10⁹ colony-forming units (CFU). In parallel, the in vitro–grown Tn5 library was further incubated to OD_600_ ≈ 2, and 2 mL of culture were harvested and stored at −80 °C as an input reference sample. Three additional independent input replicates were generated using the same procedure for downstream TraDIS analysis.

At day 2 post-infection (acute phase), piglets were humanely euthanized and necropsied. Ileum and MLN samples were aseptically collected. Clinical signs including fever, diarrhea, and lethargy were monitored daily throughout the experiment. All procedures were conducted in compliance with Good Experimental Practices and were approved by the Ethical and Animal Welfare Committee of the University of León (Spain). For each animal, two ileum segments (2 cm each) and two sets of mesenteric lymph nodes were collected. Tissues destined for histological analysis were fixed in formalin. For recovery of output mutant libraries, tissue samples were homogenized using a high-shear homogenizer and permeabilized with 0.1% Triton X-100 in PBS for 10 min to release intracellular bacteria. Tissue homogenates from each animal were pooled and cultured in 50 mL of LB broth supplemented with kanamycin, ampicillin and chloramphenicol at 37 °C with shaking until OD_600_ ≈ 2. After overnight incubation, cultures were filtered through a 100 µm cell strainer to remove residual tissue debris. Bacterial cells were collected by centrifugation (2 mL per sample) and pellets were stored at −80 °C for genomic DNA extraction. A total of five output libraries from ileum and five from mesenteric lymph nodes were recovered.

### *Ex vivo* infection of porcine neutrophils

Porcine neutrophils were isolated from peripheral blood obtained from a single healthy donor pig. Blood was collected from the ophthalmic venous sinus by licensed veterinarians at the Experimental Animal Service (SAEx) of the University of Córdoba. Neutrophil isolation was performed by density gradient separation, yielding approximately 4 × 10^8^ neutrophils from 70–90 mL of whole blood. Isolated neutrophils were seeded at 2.5 × 10^7^ cells per T75 flask and rested overnight in RPMI supplemented with 10% (v/v) fetal bovine serum (FBS) and 2% (v/v) L-glutamine at 37 °C. Neutrophil purity was routinely >90% as assessed by light microscopy.

Following stabilization, three parallel technical replicates, each constituted by 2.5× 10^7^ cells, were infected independently with the activated Tn5 mutant library at a multiplicity of infection (MOI) of 25 bacteria per neutrophil. Infections were performed in sterile 50 mL conical tubes at 37 °C with shaking for 1 h. After infection, neutrophils were washed using PBS supplemented with gentamicin (50 µg/mL) and incubated for an additional 6 h in RPMI containing 10% FBS, 2% L-glutamine, and gentamicin.

Neutrophils were subsequently pelleted by centrifugation and permeabilized using 0.1% Triton X-100 in PBS for 10 min to release intracellular bacteria. Triton X-100 was removed by centrifugation, and bacterial pellets were pooled and resuspended in LB broth supplemented with kanamycin. Cultures were incubated at 37 °C with agitation until OD_600_ ≈ 2. Then, 2 ml of the output culture were pelleted and stored for genomic DNA extraction. This pooled bacterial population constituted the neutrophil infection output library.

### Immunohistochemistry of the *in vivo* infected tissue

To prioritize samples with clear evidence of bacterial colonization, ileum and MLN sections from each animal were evaluated by immunohistochemistry using a *Salmonella*-specific antibody. Tissue samples were embedded in paraffin, sectioned at 5 μm, and mounted on Poly-L-lysine–coated slides (Sigma-Aldrich, St. Louis, MO, USA). Sections were dried overnight at 37 °C, deparaffinized in xylene, and rehydrated through graded alcohol to distilled water.

For immunohistochemistry, antigen retrieval was performed by heat treatment in 0.01 M citric acid buffer. Sections were incubated overnight at 4 °C with a rabbit polyclonal anti-*Salmonella* antibody (AgH, 1:200), followed by a 1 h incubation at 37 °C with a biotinylated anti-rabbit Ig secondary antibody (Dako, Barcelona, Spain). Antibody binding was visualized using a standard avidin–biotin peroxidase system (Dako, Barcelona, Spain) with diaminobenzidine (DAB; Sigma-Aldrich, St. Louis, MO, USA) as chromogenic substrate. To assess specificity, negative controls omitting the primary antibody were included in each assay. After development, sections were counterstained with hematoxylin, dehydrated, and mounted.

Slides were examined using an Olympus BX43 microscope equipped with an Olympus XC50 digital camera. For each infected animal, ten images were acquired from ileal sections for quantitative analysis. MLN sections were evaluated qualitatively due to lower bacterial burden. Digital image analysis was performed using Fiji (v2.17.0) with the colour deconvolution plugin to isolate the DAB signal. A fixed intensity threshold was applied uniformly to all images within the dataset to quantify the percentage of DAB-positive area.

### gDNA extraction, Tn5 amplicon library preparation and Illumina sequencing

Genomic DNA (gDNA) was extracted from the selected Tn5 mutant library pellets (4 control/input and 7 output condition samples) using the PureLink™ Genomic DNA Mini Kit (Invitrogen), following the manufacturer’s protocol. For Illumina sequencing of Tn5 insertion sites, amplicon libraries were generated for each Tn5 library following the protocol described by Karash *et* al. ^17^. Briefly, a Tn5-specific dual priming oligonucleotide (Tn5-DPO; see Supplementary Table 1) was used to perform linear PCR extension from the genomic junctions adjacent to the transposon. Purified extension products were subsequently poly-C–tailed using terminal deoxynucleotidyl transferase (TdT; New England Biolabs) and then amplified by exponential PCR using primers designed to introduce Illumina-compatible sequencing adapters and indices. Given that sequencing was performed on a NovaSeq 6000 platform, Nextera-type adapters and dual 10-bp index sequences were incorporated to allow multiplexed sample demultiplexing. A full list of primer sequences is provided in the Supplementary Table 1. Final PCR products were separated by agarose gel electrophoresis, and DNA fragments in the size range of 300–600 bp were excised and purified using the FavorPrep™ Gel/PCR Purification Kit (Favorgen).

The sequencing libraries were sequenced at the Genomics Unit of Centro Andaluz de Biología Molecular y Medicina Regenerativa (CABIMER). Each library underwent quality control, including DNA quantification by Qubit fluorometry and fragment size assessment using a 2100 Bioanalyzer (Agilent). Additionally, samples were repurified to remove adapter dimers or large DNA fragments prior to sequencing. Samples were sequenced on a NovaSeq 6000 SP-300_XP flow cell using paired-end reads (2 × 150 bp), with PhiX control DNA spiked into the sequencing pool to increase sequence diversity. Preliminary data processing was performed at CABIMER and included demultiplexing based on barcode sequences, filtering by read length and quality, adapter trimming, and N-mask removal.

### TraDIS data analysis and functional enrichment analyses

For TraDIS analysis, only forward reads were used, as reverse reads showed a high density of poly-G sequences and were therefore excluded from downstream analyses. Raw reads were first trimmed using Cutadapt v5.0 at both 5’ and 3’ ends to ensure all reads started at the GACAG motif located immediately upstream of the transposon-genome junction and to remove residual poly-G stretches ^18^. Read quality was subsequently assessed using FastQC v0.12.1.

Insertion sites were identified and gene-level analyses were performed using the Bio-Tradis pipeline ^19^. Sequencing reads from each library were mapped to the Salmonella enterica serovar Typhimurium DT104 SP11 reference genome using the bacteria_tradis mapping pipeline. Resulting BAM and plot files were processed into gene-level insertion tables using the tradis_gene_insert_sites script. Gene essentiality within each library was estimated using the tradis_essentiality.R script. Comparative fitness analyses between control/input and output/condition libraries were performed using the tradis_comparison.R. For each gene in the porcine host-conditioned core genome, the standard deviation of log_2_ fold-change across ileum, lymph node and neutrophil datasets was calculated. The distribution was summarized using the median to avoid bias from high-variance outliers. Genes with a log_2_ fold change ≤ −2 and an adjusted p-value (q-value) ≤ 0.05 were defined as conditionally essential genes (CEGs), while genes with log_2_ fold change ≥ 2 and q-value ≤ 0.05 were considered fitness-gain mutants.

Fitness comparison data were further used for data visualization using GraphPad Prism 9 and for functional enrichment analyses using the STRING database (version 12.0). Functional terms discussed in the main text were selected based on a combination of statistical significance (FDR ≤ 0.05), enrichment strength, and signal score, while minimizing redundancy across databases.

## Results

### Generation of a high-saturation transposon mutant library

To systematically define the genetic determinants supporting *S.* Typhimurium fitness across distinct porcine infection niches, we generated a highly saturated mini-Tn5 transposon mutant library in the multidrug-resistant DT104 SP11 strain using the pBAM1 delivery system. Four independent conjugations yielded a pooled library comprising approximately 1.2 million independent mutants (**Figure 1A**), providing dense genome-wide coverage suitable for high-resolution fitness profiling.

Because the pBAM1 system can generate kanamycin-resistant pseudomutants retaining the delivery plasmid without chromosomal insertion, this frequency was quantified by PCR screening of 60 randomly selected clones. The plasmid-encoded tnpA gene was detected in three isolates, corresponding to an estimated 5% pseudomutant rate (**Figure S1D**). Insertion randomness and clonal diversity were further assessed by PCR profiling of 20 mutants derived from independent conjugations, each displaying unique amplification patterns (**Figure S1F**). Sanger sequencing of five insertion junctions confirmed unique chromosomal integration sites within the DT104 SP11 reference genome (NCBI Accession CP176561.1/CP176560.1).

Collectively, these analyses demonstrate that the input library is large, genetically diverse, minimally contaminated by plasmid-retaining clones, and suitable for in vivo TraDIS screening.

### Distinct host-associated environments impose divergent selective pressures

To capture niche-specific genetic requirements during infection, the transposon library was subjected to selection in three biologically linked environments: the ileal mucosa following oral infection, primary porcine neutrophils *ex vivo*, and mesenteric lymph nodes (MLN) during systemic dissemination.

Successful intestinal invasion and systemic spread were confirmed by immunohistochemistry (**Figure 1B**). DAB-positive staining demonstrated the presence of *S*. Typhimurium within ileal villi and Peyer’s patches, and in MLN tissue, indicating dissemination beyond the intestinal barrier. Quantification of DAB-positive area in ileal samples confirmed consistent bacterial colonization across infected animals (**Figure 1C**). Based on infection reproducibility, ileum (animals 1, 2, 3) and MLN (animals 1, 2, 4) output libraries were selected for sequencing.

Across all libraries, 469.5 million reads were obtained (mean 42.7 million per sample; Supplementary Table 2). Read quality was high (mean Phred score >28), and 73.7% of reads contained Tn5-specific sequence, indicating efficient enrichment. Mapping to the DT104 SP11 genome yielded a mean alignment rate of 53.5%, with comparable insertion density across biological replicates despite variation in mapping efficiency.

Input libraries achieved near-saturation mutagenesis, with an average insertion every 36.7 bp (142,407 unique insertion sites, UIS). In contrast, output libraries displayed markedly reduced diversity (mean 39,170 UIS), reflecting selective depletion during host-associated selection **(Figure 1D**). The most pronounced contraction in insertion diversity was observed in MLN samples, consistent with a population bottleneck during systemic persistence.

Multidimensional scaling based on gene-level insertion frequencies demonstrated clear segregation between input and output libraries (**Figures 1E–G**). Output samples clustered according to anatomical niche, indicating reproducible and compartment-specific selective pressures. MLN libraries clustered furthest from the input population, supporting the notion of progressively stringent selection along the infection trajectory.

Differential abundance analysis identified genes significantly depleted during infection (log₂FC ≤ −2; q ≤ 0.05), consistent with conditional essentiality for colonization and dissemination (**Figures 1H–J**). Together, these data demonstrate that each host niche imposes a distinct selective landscape that reshapes the mutant population.

### Basal essentiality and host-conditioned fitness requirements define complementary genetic landscapes

To distinguish basal from infection-dependent genetic requirements, essential genes for growth in rich medium were defined using the four independent input libraries. Essentiality calls were highly reproducible across replicates (**Figure S2A**), identifying 438 genes consistently devoid of insertions and therefore essential under laboratory conditions (Supplementary Table 3).

The intersection of essential genes across all libraries defined the core essential genome, comprising 429 genes that represent the minimal genetic requirements for viability irrespective of environmental context. Functional enrichment analysis revealed strong overrepresentation of fundamental cellular processes, including translation (GO:0006412), ribosome biogenesis (KEGG: stm03010), and macromolecule biosynthesis (GO:0034645), consistent with the canonical structure of a bacterial essential genome. The complete gene list is provided in Supplementary Table 3, and enrichment results are detailed in Supplementary Table 4. Comparison with the essential genome previously defined in *S.* Typhimurium SL1344^20^ demonstrated substantial overlap (**Figure S2B**), supporting the robustness and completeness of the DT104 dataset while highlighting strain-specific variation.

Across all host-associated conditions examined, genes classified as essential *in vitro* remained uniformly depleted of insertions, confirming that infection-driven selection operated upon a conserved basal framework. However, beyond this core essential genome, each infection niche imposed additional layers of conditional essentiality, reflecting the distinct physiological and immunological constraints encountered by *S*. Typhimurium within the porcine host.

To resolve these host-conditioned requirements, Tn5 insertion profiles were compared between input libraries and output populations recovered from ileum, primary porcine neutrophils, and MLN. Neutrophil-derived libraries originated from a single biological donor with three parallel infection replicates, whereas ileum and MLN libraries were obtained from independent animals.

Conditionally essential genes (CEGs) were defined as loci significantly depleted during infection (log₂FC ≤ −2; q ≤ 0.05), reflecting genetic functions required to withstand host-imposed stresses such as immune pressure, nutrient restriction, and intracellular antimicrobial mechanisms. Conversely, fitness-gain mutants were defined as loci whose disruption significantly increased bacterial representation during infection, indicating functions that are disadvantageous in specific host environments and whose loss confers a selective advantage.

Comparative analysis revealed 368 CEGs and 78 fitness-gain mutants in the ileum. Neutrophil-derived samples yielded 657 CEGs and 270 fitness-gain mutants, whereas MLN samples exhibited a markedly larger set of 1,370 CEGs but only 36 fitness-gain mutants (**Figures 1H–J**; Supplementary Table 3). The progressive increase in CEGs from ileum to MLN indicates escalating selective stringency during dissemination, consistent with increasing host-imposed constraints along the infection trajectory. In contrast, the disproportionately high number of fitness-gain mutants in neutrophils suggests that adaptive loss of specific functions confers a selective advantage within this intracellular niche, potentially reflecting metabolic streamlining, modulation of immune-detectable pathways, or suppression of energetically costly processes under oxidative stress.

The degree of overlap between niches further underscores compartmental specialization. Most CEGs were condition-specific, with only a limited subset shared across all three environments (**Figure 2A**), indicating that intestinal colonization, survival within neutrophils, and MLN persistence rely on partially distinct genetic programs. Similarly, fitness-gain mutants displayed minimal cross-niche overlap (**Figure 2B**), demonstrating that adaptive loss-of-function events are highly context-dependent rather than broadly beneficial across host tissues.

**Figure 2.**
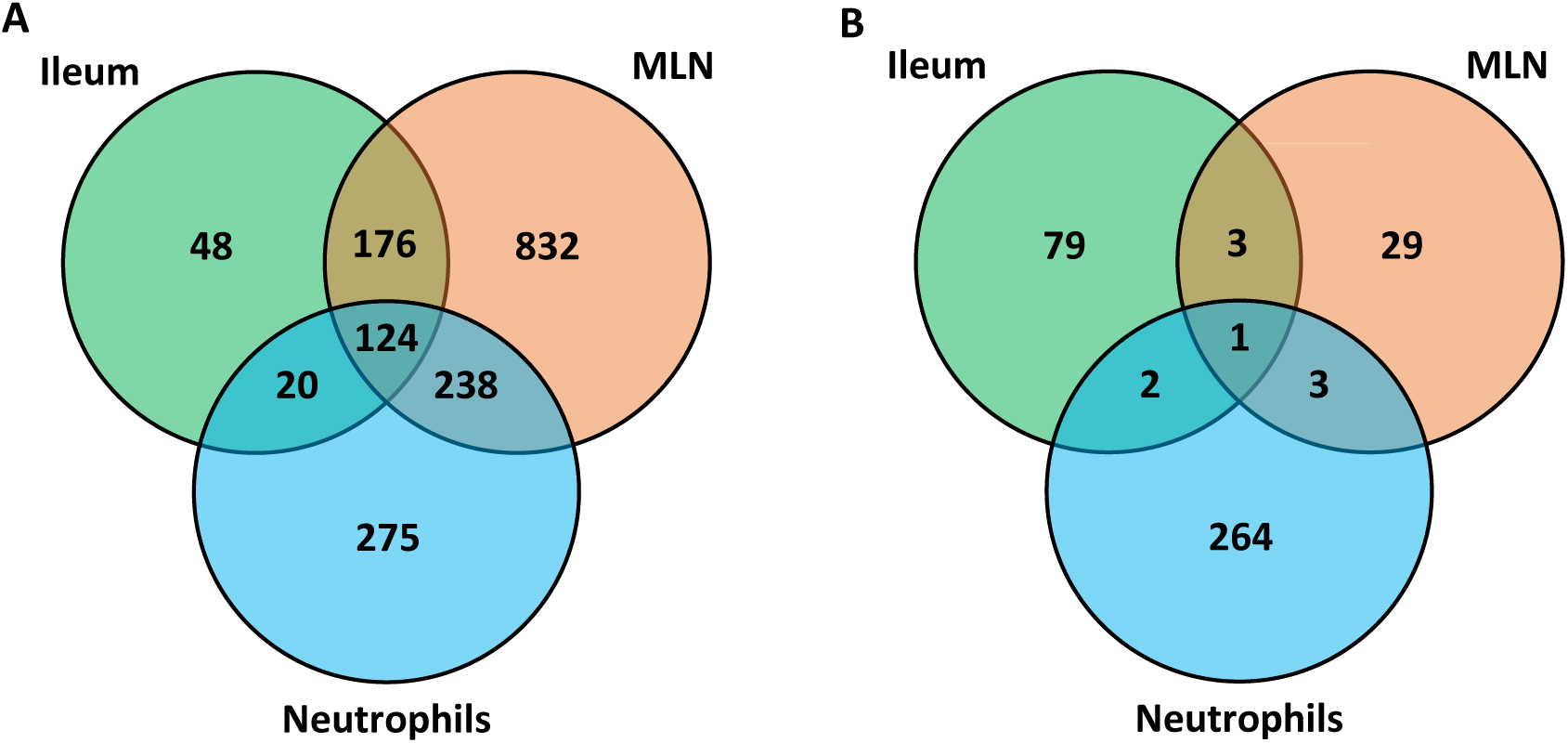
Comparative Overlap of Condition-Specific of Conditional Essential Genes (CEGs) and Fitness-Gain Mutants. **A.** Venn diagram displaying the number of CEGs identified in each of the conditions of study. CEGs were filtered out based on their log_2_FC (≤-2) and q.value (≤ 0.05). **B**. Venn diagram displaying the number of fitness-gain mutants identified in each of the conditions of study. Fitness-gain mutants were filtered out based on their log_2_FC (≤-2) and q.value (≤ 0.05).

Together, these findings define complementary genetic landscapes in which conserved basal requirements coexist with niche-specialized and host-imposed constraints. *S*. Typhimurium adaptation to porcine infection thus emerges from the interplay between a stable essential genomic backbone and dynamically reconfigured fitness programs shaped by the anatomical and immunological environment.

### Tissue-specific fitness landscapes across the porcine infection route

#### Ileal colonization selects for invasion, spatial positioning and nutrient acquisition

In the ileal tissue, a total of 368 genes *S.* Typhimurium DT104 SP11 may be needed for a successful invasion of the tissue by the pathogen. To functionally resolve these CEGs, enrichment analysis was performed using STRING. Of the 368 CEGs, 249 mapped to the STRING database and clustered into four major functional groups based on k-means similarity networks: nucleic acid binding (79 genes), virulence-associated functions (74 genes), lipopolysaccharide (LPS) biosynthesis (56 genes), and cellular amino acid biosynthesis (32 genes) (**Figure 3A**). Enrichment analysis further highlighted pathways linked to virulence and secretion (CL:5322), flagellar assembly and chemotaxis (CL:5087), O-antigen/LPS biosynthesis (GO:0033692; GO:0009243), and amino acid biosynthetic processes (CL:1815). DNA repair (GO:0006281) was also significantly enriched (FDR < 0.05), although with lower enrichment strength (Supplementary Table 4).

**Figure 3.**
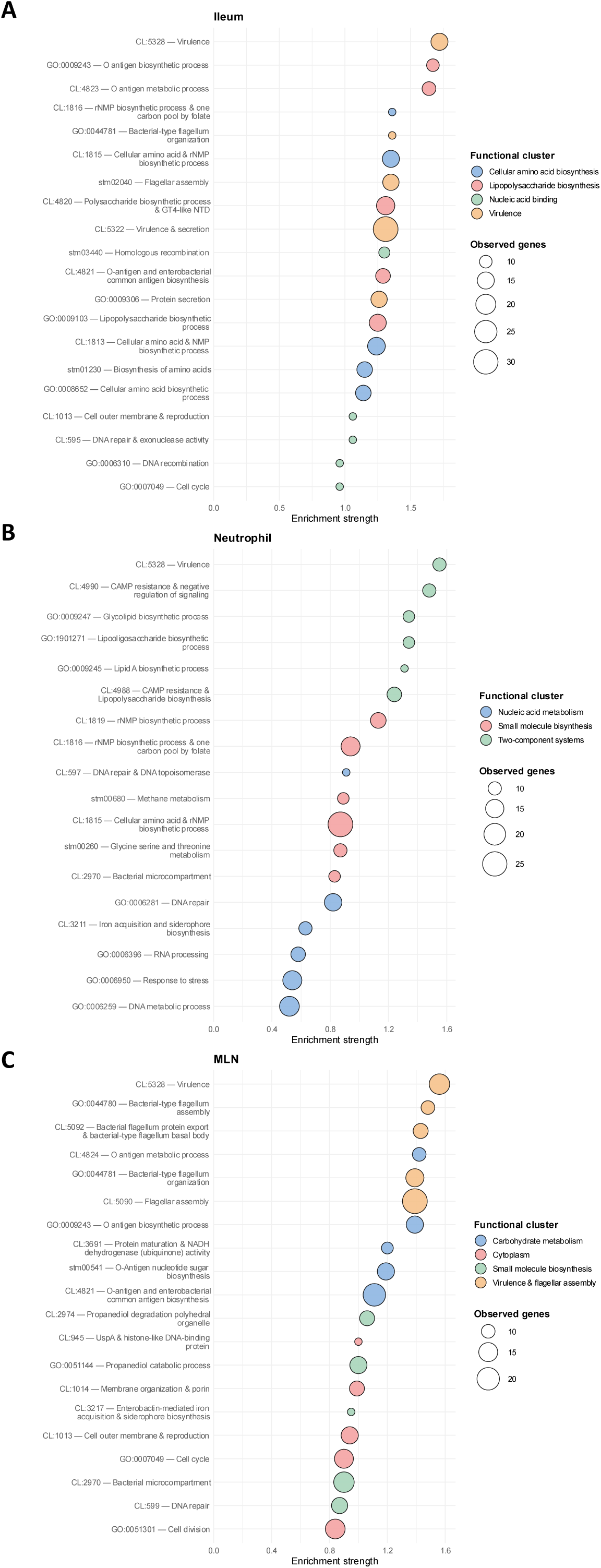
Functional enrichment of *S.* Typhimurium CEGs across porcine tissues. Core essential genes (CEGs) identified in each tissue were mapped to the STRING v12.0 database for functional enrichment analysis. Enriched terms were grouped into functional clusters using the STRING k-means clustering algorithm. For each cluster, the top five terms were selected based on the enrichment strength parameter. Bubble size represents the number of observed genes per term, and the x-axis indicates enrichment strength. Colours denote functional clusters. **A.** Ileum. **B**. Neutrophil. **C.** Mesenteric lymph nodes (MLN).

Consistent with the central role of active invasion during intestinal colonization, multiple genes encoded within SPI-1 and SPI-2 emerged as key ileum-associated determinants. Among the 48 ileum-specific CEGs, defined as genes exclusively essential in this compartment (**Figure 2A**), five virulence genes were identified (*spiA, spiC, ssaG, ssaP* and *sipB*), four of which are encoded within SPI-2 (**Figure 4A**). The detection of SPI-2 components at this early stage suggests that intracellular adaptation programs may be engaged shortly after epithelial invasion, potentially reflecting rapid uptake by epithelial cells or resident phagocytes.

**Figure 4.**
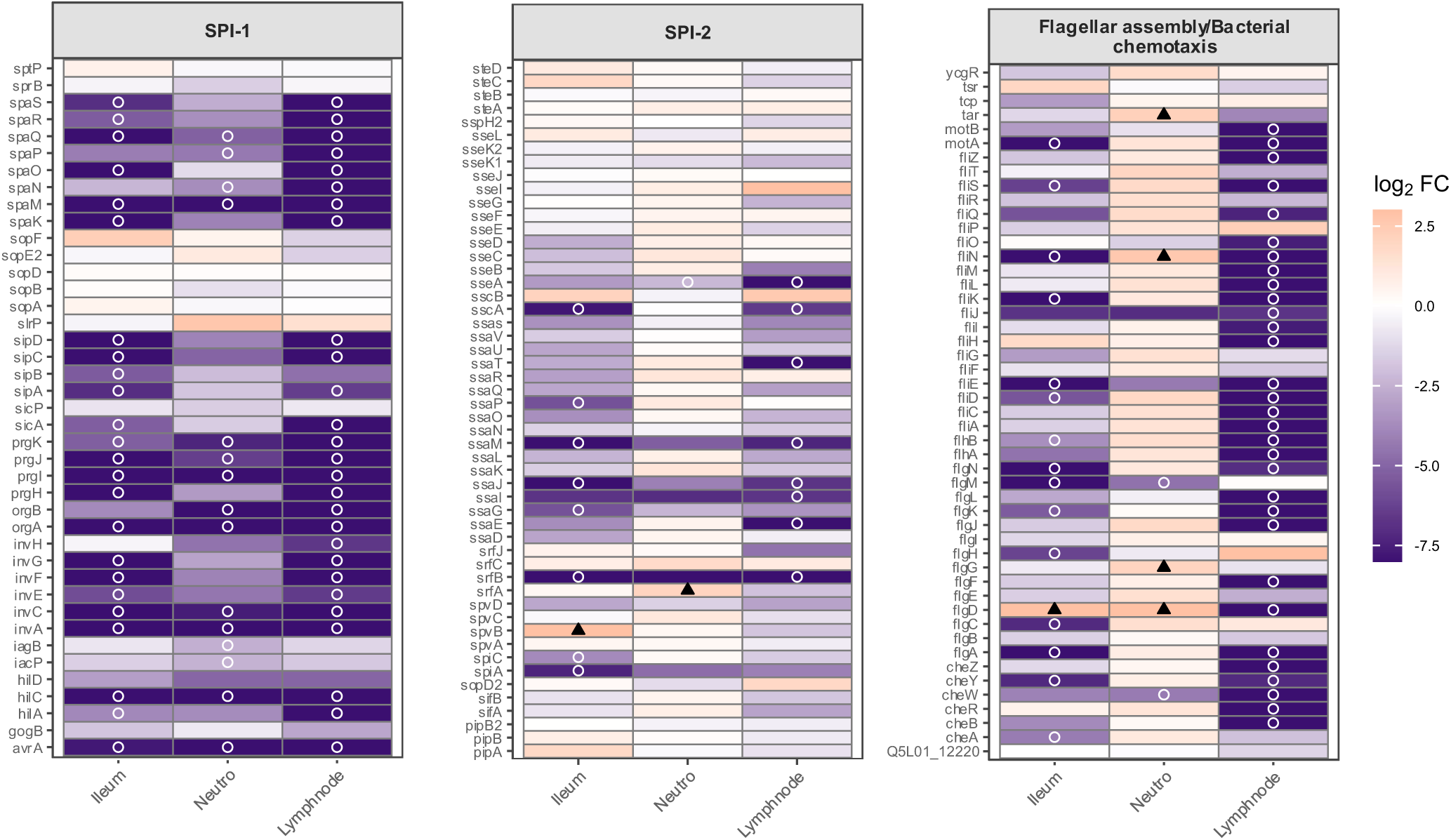
Niche-specific distribution of virulence determinants and *Salmonella* pathogenicity islands during porcine infection. Heatmap representation of log₂ fold-change values for selected virulence and regulatory genes across compartments, as informed by the TraDIS screen with q. value ≤ 0.05 marked with “circle”. **A**. Virulence and secretion (STRING cluster CL:5322). **B**. Flagellar assembly and bacterial chemotaxis (STRING cluster CL:5089).

Beyond secretion systems, ileal fitness was strongly influenced by bacterial motility and chemotaxis. Four ileum-specific CEGs (*flgC, flgH, flhD, cheA*) are directly involved in flagellar biosynthesis and directional movement, underscoring the importance of spatial positioning within the complex architecture of the intestinal mucosa (**Figure 4B**). Efficient navigation likely facilitates epithelial contact, microenvironmental sensing, and access to nutrient-rich niches. Envelope integrity also emerged as a critical determinant. The enrichment of LPS and O-antigen biosynthetic genes indicates that maintenance of outer membrane robustness is required to withstand host-derived antimicrobial peptides and inflammatory stress (**Figure 5**).

**Figure 5.**
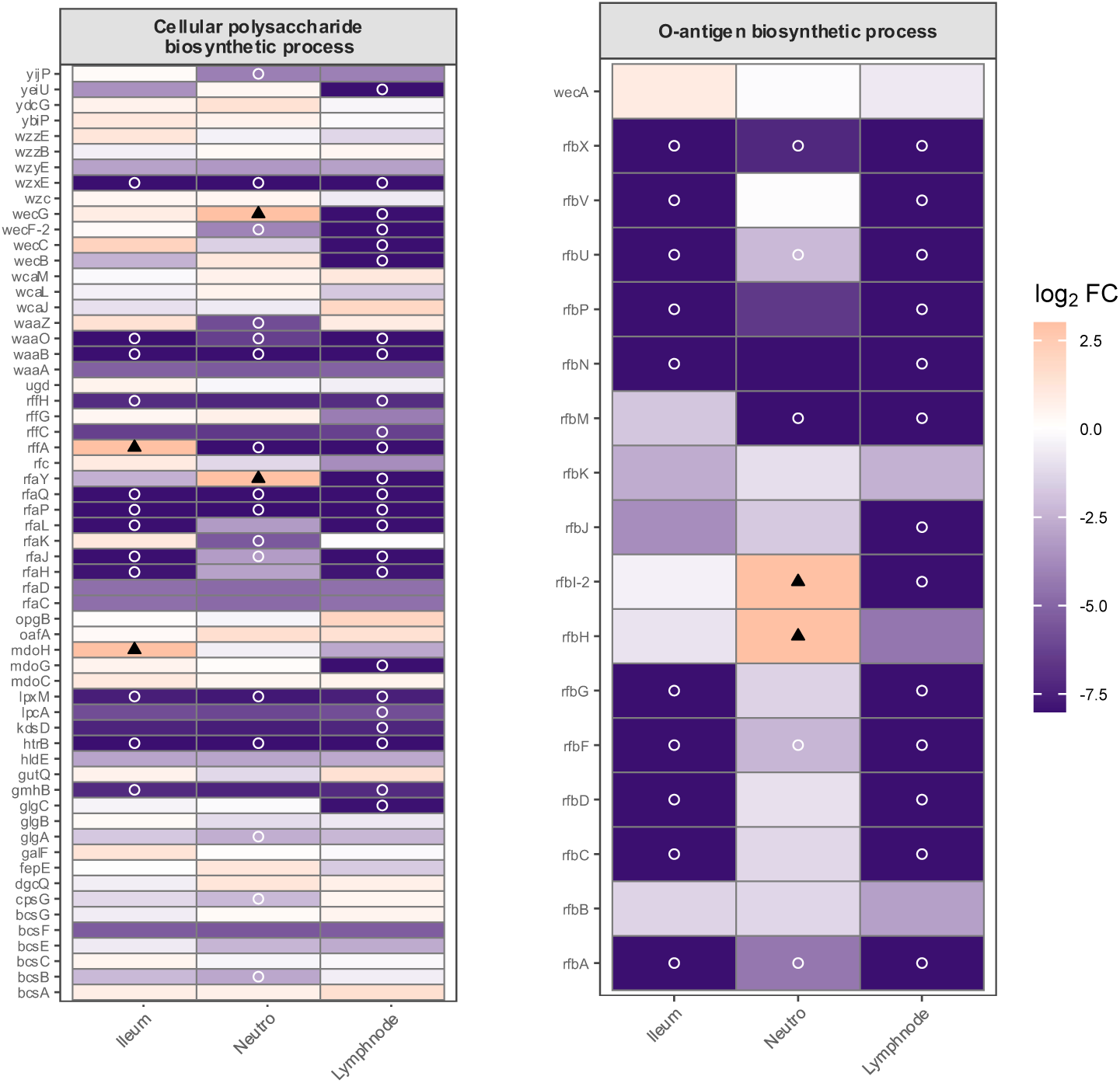
LPS biosynthesis determinants during *Salmonella* infection in porcine hosts. Heatmap representation of log₂ fold-change values for selected cellular polysaccharide biosynthesis (GO:0033692) genes across compartments, as informed by the TraDIS screen with q. value ≤ 0.05 marked with “circle”.

Metabolic adaptation represented an additional layer of ileal specialization. Five ileum-specific CEGs were associated with nutrient acquisition and transport systems (t*hiQ, Q5L01_23530, Q5L01_23990, Q5L01_18335, deoD;* Supplementary Table 3), while broader enrichment of amino acid biosynthetic pathways suggests that nutrient limitation within the porcine ileum imposes biosynthetic demands on the pathogen. These findings are consistent with competition against the resident microbiota and the need to exploit host- and diet-derived substrates during early colonization.

Collectively, functional clustering and niche-specific gene analysis converge on a model in which the ileum selects for *S*. Typhimurium variants capable of coordinated deployment of invasion machinery, precise spatial navigation, envelope resilience, and metabolic flexibility.

### Survival within porcine neutrophils is driven by stress resistance and metabolic rewiring rather than classical virulence

Selection of the transposon mutant library within primary porcine neutrophils generated a fitness landscape strikingly distinct from that observed in the intestinal mucosa. Using an *ex vivo* infection model based on neutrophils isolated from a single donor pig with three parallel technical replicates, we identified 657 CEGs, of which 382 mapped to the STRING database. K-means clustering resolved three major functional networks: small-molecule biosynthesis (163 genes), nucleic acid metabolism (119 genes), and two-component regulatory systems (59 genes) (**Figure 3B**). Functional enrichment analysis revealed three dominant biological themes: virulence-associated functions (CL:5328), stress response pathways (CL:4990; stm02020; GO:0006281), and secondary metabolic rewiring (stm01110; GO:0008652; CL:2969) (Supplementary Table 4). However, in contrast to the ileal niche, classical invasion-associated systems contributed minimally to neutrophil fitness.

A total of 275 neutrophil-specific CEGs were identified **(Figure 2A**), predominantly associated with resistance to cationic antimicrobial peptides (CAMP) (*arn* operon*, pagP*), oxidative and envelope stress responses (*cpxA, oxyR, uvrY*), and metabolic adaptation pathways (*pduGQ, eutHL*) (**Figure 6**; Supplementary Table 3). Genes associated with amino acid biosynthesis, cofactor production, anaplerotic reactions, and alternative substrate utilization were selectively required, indicating that intracellular persistence demands extensive metabolic reprogramming. Notably, bacterial microcompartment-associated operons *(pdu* and *eut*) were identified as conditionally essential, suggesting that *S*. Typhimurium exploits specialized metabolic modules to sustain viability under nutrient-restricted phagolysosomal conditions. In contrast, only a limited number of virulence-related determinants were uniquely required in this compartment (**Figure 4**).

**Figure 6.**
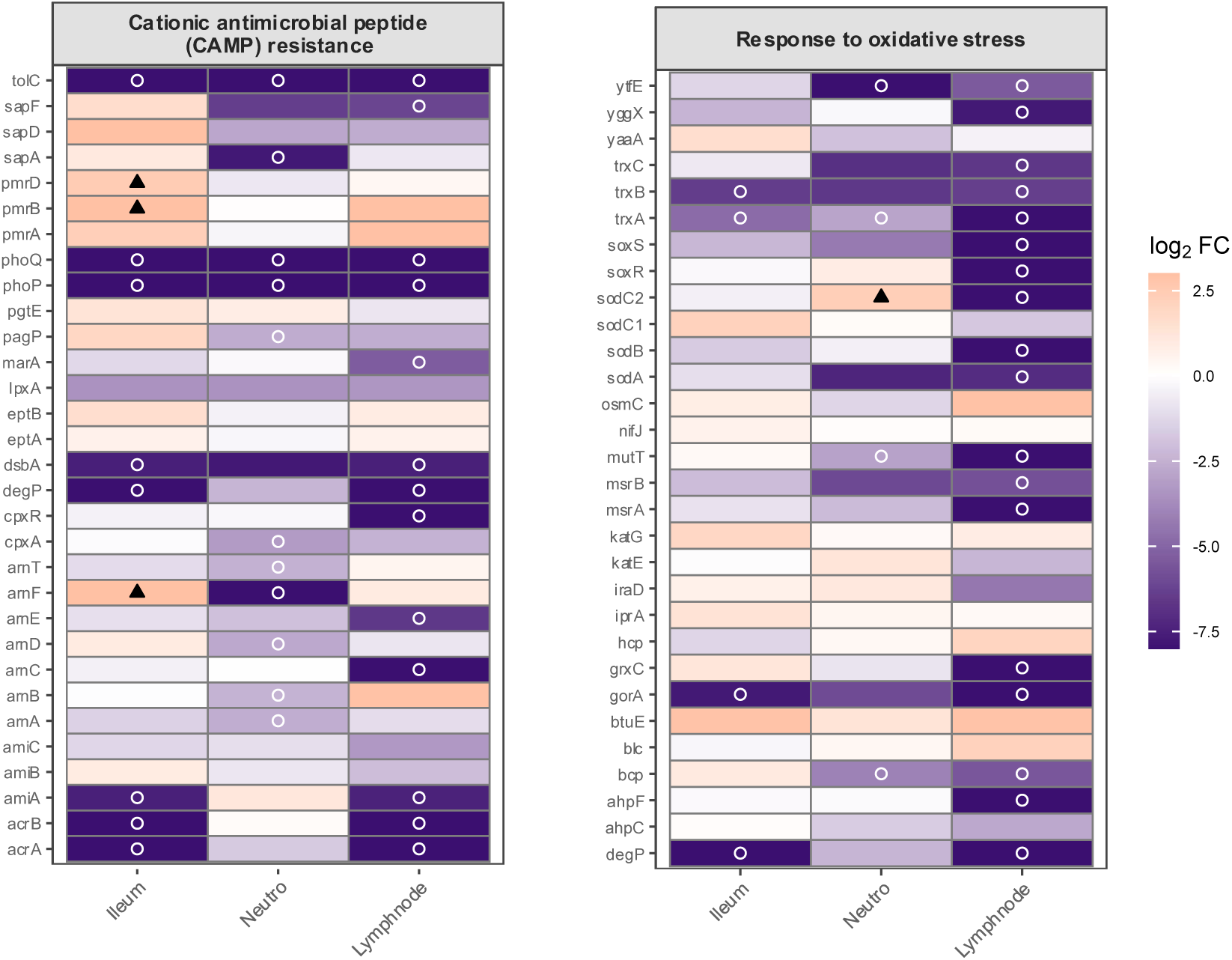
Antimicrobial peptide resistance determinants and oxidative stress defence during *Salmonella* infection in porcine hosts. Heatmap representation of log₂ fold-change values of each gene across infection conditions; genes with q.value ≤ 0.05 are marked with “circle”. **A**. Cationic antimicrobial peptide (CAMP) resistance (KEGG stm01503). **B**. Response to oxidative stress (GO:0006979).

Strikingly, neutrophil selection was also characterized by a disproportionately high number of fitness-gain mutants relative to other niches (**Figure 1H–J**). These advantageous disruptions were enriched among genes involved in surface structures, regulatory systems, and metabolic functions, indicating that loss of certain bacterial activities may reduce immune detection or alleviate metabolic burden under intracellular stress. This pattern suggests that survival within neutrophils may favor streamlined physiological states rather than maximal virulence expression.

Collectively, these findings define porcine neutrophils as a niche that imposes stringent oxidative and nutritional constraints, selecting for *S*. Typhimurium variants optimized for stress resistance and metabolic economy. This survival strategy contrasts sharply with the invasion- and motility-focused requirements observed in the ileum, highlighting the dynamic reconfiguration of bacterial fitness landscapes as the pathogen transitions between host environments.

#### Persistence within mesenteric lymph nodes reveals the most restrictive host-imposed fitness landscape

Analysis of transposon mutant representation in bacteria recovered from MLN revealed the most stringent selective environment encountered by *S*. Typhimurium along the porcine infection route. MLN samples displayed the largest contraction in mutant diversity and the highest number of CEGs (n = 1,370), consistent with a pronounced population bottleneck during systemic persistence.

Of these CEGs, 701 mapped to the STRING database and clustered into four principal functional networks: small-molecule biosynthesis (266 genes), cytoplasmic functions (163 genes), carbohydrate metabolism (140 genes), and virulence/flagellar assembly (122 genes) (**Figure 3C**). Functional enrichment analysis revealed strong dependence on pathways related to motility and chemotaxis (CL:5089; GO:0097588), virulence and secretion systems (CL:5322), lipopolysaccharide biosynthesis (GO:0033692; GO:0009243), and extensive metabolic reprogramming (stm01110; CL:1815; CL:2969; GO:0044283). In contrast, classical stress-response pathways (GO:0006281; GO:0033554; stm02020) and cell cycle functions (CL:1013; GO:0007049) showed lower enrichment strength and higher FDR values, suggesting a comparatively secondary contribution (Supplementary Table 4).

Remarkably, 832 genes (60%) were tissue-specific CEGs in the MLN (**Figure 2A**), underscoring the exceptional selectivity of this compartment. Among these were nine genes belonging to three major secretion systems (**Figure 4**): components of the SPI-1 type III secretion system, SPI-2–encoded type III secretion machinery, and the type VI secretion system (*Q5L01_19415, tssG, Q5L01_19435*), the latter associated with interbacterial competition. The simultaneous requirement for SPI-1, SPI-2 and T6SS components suggests that persistence within organized lymphoid tissue demands coordinated deployment of multiple virulence strategies.

Motility and chemotaxis genes were also prominently represented among MLN-specific determinants, including multiple components of the flagellar apparatus and chemotaxis regulators (**Figure 4**). This requirement implies that active positioning within lymphoid microenvironments may facilitate access to permissive intracellular niches or escape from hostile immune microdomains.

Metabolic adaptation constituted a major fraction of MLN-associated requirements. Genes associated with bacterial microcompartments (*pdu* operon*, eutK, eutA*), secondary metabolism, and alternative respiration pathways were selectively required. In particular, genes involved in the vitamin K/menaquinone cycle (*menEFH, ubiI*), essential for anaerobic respiration, and the enterobactin-mediated iron acquisition system (*ent* operon) were conditionally essential, consistent with nutritional immunity and oxygen limitation within lymphoid tissue (Supplementary Table 3).

Moreover, *S.* Typhimurium employs a set of 22 genes specialized in stress response, including oxidative, acid, envelope and general stress systems (*uspA, cpxR*, *pspF*; Supplementary Table 3), indicating that sustained immune pressure still contributes to selection in this niche.

These findings portray the MLN as the most restrictive environment encountered during porcine infection, requiring coordinated engagement of virulence systems, motility, envelope integrity, iron acquisition, anaerobic respiration, and extensive metabolic flexibility. In contrast to the invasion-focused profile of the ileum and the stress-resistance strategy dominant in neutrophils, MLN persistence demands an integrated and comprehensive genetic program.

### Cross-niche convergence reveals a conserved porcine host-conditioned essential genome

Despite the pronounced specialization of *S*. Typhimurium fitness landscapes across the ileum, neutrophils and MLN, a defined subset of genes was consistently required in all three host-associated environments. A total of 124 CEGs were strictly conserved across infection sites (**Figure 2A**), defining a core porcine host-conditioned essential genome. These genes were dispensable for *in vitro* growth but reproducibly depleted in every *in vivo* and *ex vivo* host condition, indicating that they represent fundamental adaptive requirements imposed by the porcine host rather than niche-restricted functions. Their recurrence across anatomically and physiologically distinct tissues supports strong and consistent selective constraints acting throughout the infection trajectory.

Functionally, this conserved gene set delineates five coherent biological axes that underpin *in vivo* fitness: (i) a minimal SPI-1-encoded T3SS invasion backbone, (ii) a conserved module of outer membrane integrity factors, (iii) post-transcriptional regulation and RNA stability, (iv) genome repair and maintenance, and (v) a metabolic backbone encompassing central carbon metabolism, amino acid biosynthesis, the shikimate pathway and cofactor biosynthesis.

Notably, two systems emerged as universally indispensable across all host environments analyzed: the Mla pathway, which maintains outer membrane lipid asymmetry^21,22^ and the twin-arginine translocation (Tat) system, responsible for the export of folded proteins to the periplasm^23^. Disruption of either pathway resulted in severe fitness defects in every infection niche examined, indicating that maintenance of envelope integrity and efficient protein trafficking constitute fundamental requirements for *S*. Typhimurium survival *in vivo.* Their consistent necessity across ileal, neutrophil and MLN environments, despite distinct immune and metabolic pressures, underscores their role as core determinants of bacterial viability and host adaptation.

Together, these findings support a hierarchical model of *S*. Typhimurium host adaptation in pigs, in which niche-specific genetic programs operate atop a shared set of host-imposed physiological constraints. By delineating both dimensions of bacterial fitness, this study provides a unified framework for understanding *Salmonella* persistence in a large-animal host and identifies conserved pathways that may represent attractive targets for intervention strategies aimed at controlling non-typhoidal salmonellosis at the animal–human interface.

## Discussion

In this study, we used a genome-wide TraDIS approach to define the genetic determinants of *Salmonella enterica* serovar Typhimurium fitness across sequential host niches during infection of pigs, a highly relevant large-animal reservoir for zoonotic transmission. By integrating *in vivo* and *ex vivo* selection in the ileal mucosa, primary porcine neutrophils and MLN, we provide a comprehensive view of how bacterial genetic requirements are dynamically reshaped as Salmonella progresses along its natural infection route within a single host (**Figure 7**). Given the close resemblance between porcine and human non-typhoidal salmonellosis, this model enhances the translational value of *in vivo* transposon profiling.

**Figure 7.**
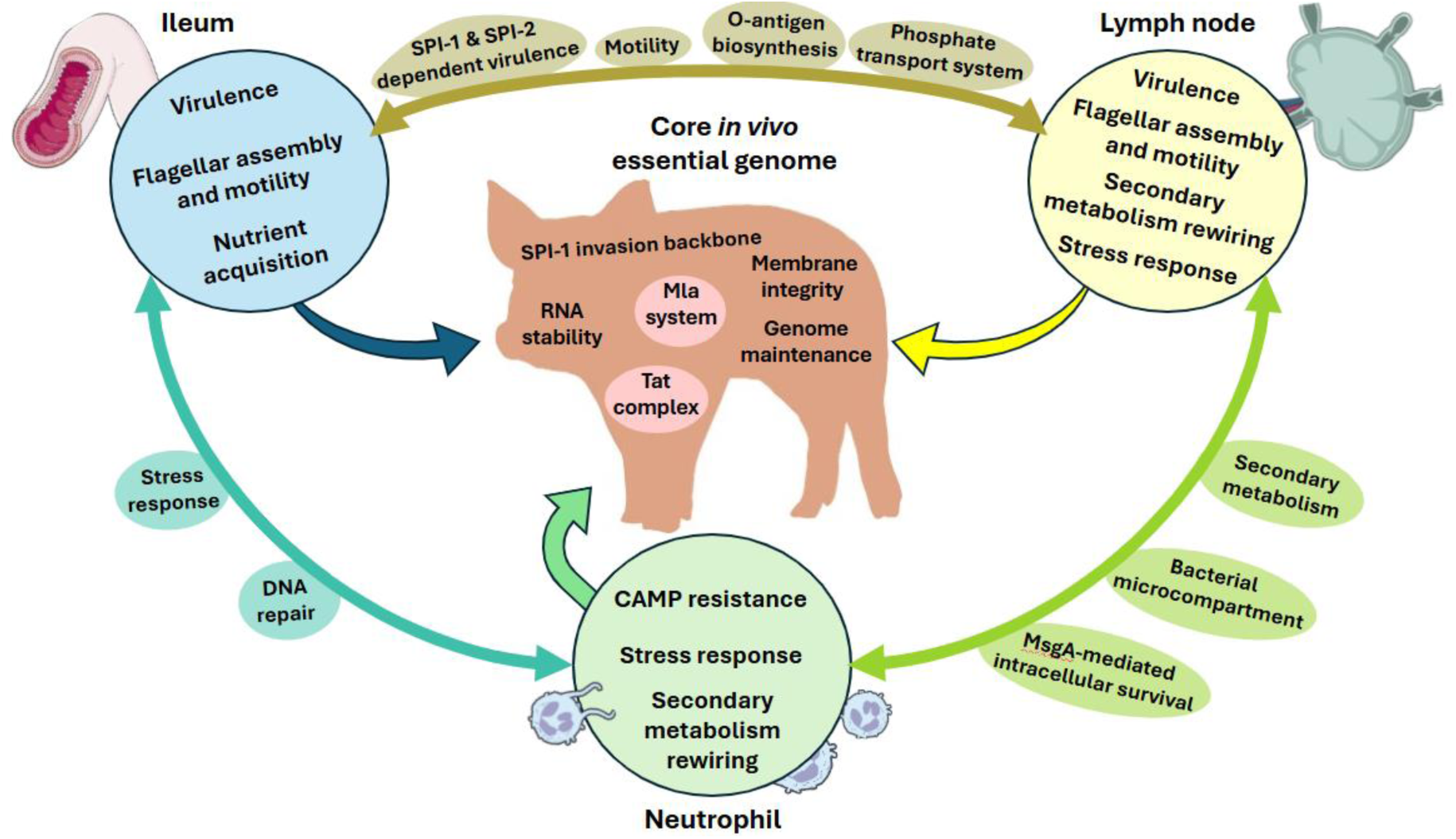
Tissue-specific fitness landscapes and conserved in vivo essential genome of *S*. Typhimurium in the porcine host. Schematic representation summarizing niche-imposed genetic constraints during in vivo infection of porcine ileum, lymph node and neutrophils.

Previous genome-scale fitness studies of *Salmonella* have largely focused on single anatomical compartments or relied on parenteral murine models, limiting insight into how selective pressures evolve across the natural course of infection. Although TraDIS analyses have been applied to food-producing animals after oral challenge, these studies primarily interrogated isolated intestinal niches^12^. Our results extend this work by demonstrating that *S*. Typhimurium experiences profoundly distinct selective landscapes as it transitions from the intestinal mucosa to innate immune cells and organized lymphoid tissue within the same porcine host.

The progressive reduction in mutant diversity observed across niches, culminating in a pronounced bottleneck within MLN, underscores the increasing stringency of host-imposed selection during systemic persistence. This finding highlights the importance of considering infection as a dynamic, multi-stage process rather than a series of independent environments, particularly when seeking to identify determinants of long-term persistence and transmission.

Methodologically, comparison with the porcine TraDIS dataset of Chaudhuri et al.^12^ revealed modest gene-level overlap (18%) but strong convergence at the pathway level, including SPI-1 components, Tat secretion, LPS biosynthesis, and envelope-associated determinants. Importantly, both studies used pre-infection mutant libraries, thereby capturing cumulative fitness costs associated with host entry and dissemination, an aspect not addressed by post-colonization mutagenesis approaches^24^.

Our analysis reveals that each infection niche selects for a characteristic genetic program reflecting its dominant physiological constraints. In the ileum, fitness was primarily driven by genes involved in epithelial invasion, motility and nutrient acquisition, consistent with early colonization requirements. The identification of SPI-2–associated genes as ileal fitness determinants suggests that intracellular adaptation programs may be engaged earlier during porcine infection than traditionally appreciated^25^.

Neutrophil survival, in contrast, was dominated by oxidative and envelope stress resistance coupled with metabolic rewiring. Major virulence systems were comparatively dispensable, while fitness-gain mutants were enriched, indicating that intracellular persistence in neutrophils favours physiological resilience and metabolic economy over-active host manipulation. The selective requirement for coordinated *arn* operon activity^26^ supports adaptation to intense antimicrobial peptide pressure^27–30^, while repression of flagellar functions is consistent with avoidance of innate immune recognition and NET induction ^31–33^.

Persistence within MLN imposed the most comprehensive genetic program, integrating virulence-associated systems, chemotaxis, envelope integrity and broad metabolic capacity. SPI-1 and SPI-2 components displayed hierarchical and niche-dependent contributions, consistent with effector redundancy and selective deployment *in vivo*^34–36^. Extensive selection across the *fli* operon and chemotaxis genes indicates that directed motility contributes to spatial positioning within organized lymphoid tissue. Concurrently, strong requirements for iron acquisition, menaquinone (vitamin K) biosynthesis and bacterial microcompartment systems (*eut* and *pdu*) underscore the metabolic demands of sustained intracellular survival stress^37–45^. Collectively, these features position MLN as the most restrictive environment encountered during porcine infection.

Despite pronounced tissue-specific specialization, a compact set of genes emerged as universally required across all host-associated environments, defining a conserved porcine host-conditioned essential genome. This core was enriched for outer membrane integrity, genome maintenance and basal SPI-1 functions, reflecting invariant physiological constraints imposed by the host.

Strikingly, two complete systems, the Mla phospholipid transport pathway and the twin-arginine translocation (Tat) machinery, were essential across all niches. The Mla system (*mlaACDEF*) maintains outer membrane lipid asymmetry and barrier function. Although previously associated with antimicrobial resistance and permeability defects^21^, its comprehensive requirement during large-animal infection has not been demonstrated. The concurrent identification of PbgA/YejM further reinforces membrane lipid homeostasis as a central *in vivo* constraint constraint ^22,46^.

Similarly, three Tat components (*tatB, tatC* and *tatE*) were universally required. The Tat pathway exports folded proteins involved in stress tolerance and virulence^47^ and has been implicated in murine and avian models^23,48,49^. Our findings extend these observations to the porcine host and establish Tat-mediated protein trafficking as a non-negotiable requirement for systemic persistence. These findings emphasize the value of cross-niche analyses for identifying conserved vulnerabilities that may be obscured in single-environment studies.

The porcine host closely mirrors key aspects of human non-typhoidal salmonellosis, including gastrointestinal physiology, immune architecture and infection outcome. By defining *Salmonella* fitness landscapes in pigs, our study provides insights with enhanced translational relevance compared to murine-only models. The identification of conserved host-conditioned essential pathways suggests potential intervention targets aimed at limiting bacterial persistence in animal reservoirs and reducing zoonotic transmission.

More broadly, this work illustrates how large-animal infection models combined with genome-scale functional genomics refine understanding of pathogen adaptation in physiologically relevant contexts. Extending similar multi-niche approaches to other zoonotic pathogens may reveal conserved host-imposed constraints exploitable for broad-spectrum control strategies.

### Limitations and future directions

Several limitations of this study warrant consideration. Neutrophil selection was performed using cells derived from a single donor, which may limit the generalizability of specific fitness effects. However, the strong functional coherence of the identified pathways and their convergence with *in vivo* findings support the robustness of the observed trends. In addition, while TraDIS provides powerful population-level insights into fitness requirements, it does not capture temporal dynamics or cell-type–specific interactions within complex tissues. Future studies integrating single-cell approaches, temporal sampling and targeted mutant validation will further refine the genetic architecture of *Salmonella* infection in pigs.

## Author contributions

Conceptualization: S.Z.L., T.G.G. and J.J.G. Methodology: A.R.G., J.B.B. and L.B. Investigation: All authors. Writing original draft: A.R.G and T.G.G. Manuscript draft revisions and editing: J.J.G. Review and approval of draft: all authors. Funding acquisition: S.Z.L and J.J.G.

## Acknowledgements

The authors wish to express their gratitude to Héctor Arguello and Ana Carvajal (Department of Animal Health, Faculty of Veterinary Medicine, University of León, Spain) for their valuable support during the *in vivo* infection experiments.

## Supplementary Figures

**Figure S1.**
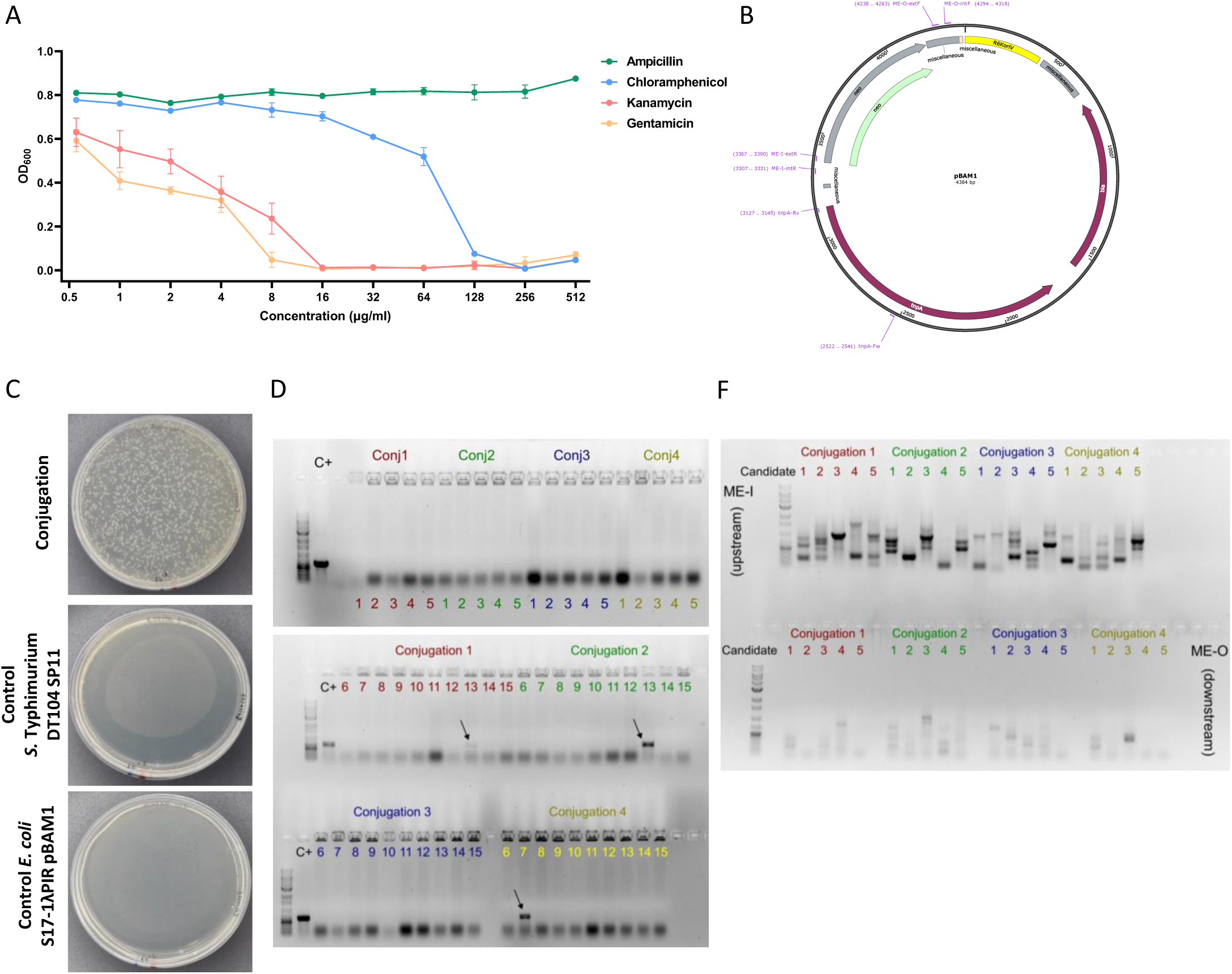
Tn5 library construction. **A.** Antibiotic susceptibility profiling of S. Typhimurium DT104 SP11 by MIC assay for ampicillin, chloramphenicol, kanamycin, and gentamicin. Data represent biological triplicates (mean ± SD). **B**. The genomic map of the pBAM1 plasmid is shown, indicating the antibiotic resistance markers (bla and neo) and the primer binding sites used for validation. **C.** Assessment of conjugation efficiency and selective conditions. Representative plates showing selective growth of S. Typhimurium DT104 SP11 Tn5 transconjugants under kanamycin and chloramphenicol selection. **D.** Agarose gel electrophoresis of colony PCR products targeting the *tnpA* gene in 60 randomly selected mutants from four independent conjugation experiments. The pBAM1 plasmid was used as a positive control (C+). Positive amplicons are indicated by an arrow. **F.** Agarose gel electrophoresis of semirandom colony PCR products amplifying the upstream (ME-I) and downstream (ME-O) genomic junctions of the Tn5 transposon in 20 representative mutants. Distinct banding patterns indicate independent chromosomal insertion events across conjugations.

**Figure S2.**
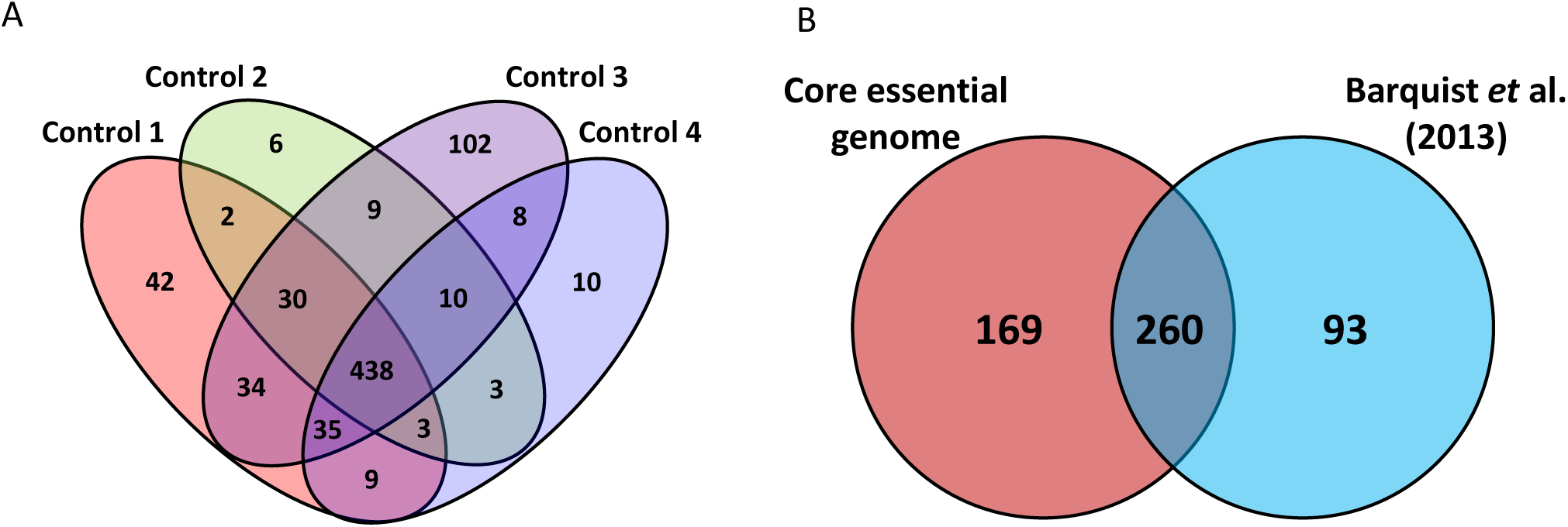
Core Essential Genome Identification and Cross-Study Comparison. **A.** Venn diagram showing the intersection of genes predicted to be essential in four independently generated Tn5 input libraries. Essentiality was inferred using the tradis_essentiality.R script from the Bio-Tradis pipeline, based on the absence or strong depletion of transposon insertions across gene bodies. The high overlap of essential genes among replicates demonstrates the robustness and reproducibility of essentiality calls across independently generated libraries. **B**. Venn diagram comparing the genes identified as the core essential genome in this study on *S.* Typhimurium DT104 SP11 and the core essential genome described in *S.* Typhimurium SL1344 by Barquist *et al*., 2013.

